# Syndecan-4 is the Key Proteoglycan Mediating LPS-induced Sepsis-associated Lung Injury

**DOI:** 10.1101/2023.05.02.539148

**Authors:** Zhipeng Zhu, Xiaoyan Ling, Hongmei Zhou, Junran Xie

## Abstract

The occurrence of acute sepsis-associated lung injury (ALI) is closely related to vascular endothelial cell dysfunction, but the exact isoform of the functional syndecans (SDCs) located in the inner endothelial membrane of blood vessels and involved in regulating inflammation is not clear. The present study aimed to clarify whether SDC2 and/or 4 is involved in the inflammation process.Using the Gene Expression Omnibus (GEO) database, differentially expressed genes (DEGs) were extracted and analyzed. A bioinformatic analysis was conducted using R language, gene ontology (GO), and the Kyoto Encyclopedia of Genes and Genomes (KEGG) with the aim of predicting SDC2 and 4 expression levels. Subsequently, the expression profiles of these two proteins were examined after construction of a sepsis-associated ALI mmodel was constructed in both human lung vascular endothelial cells (HLVEC) and animal models under inflammatory conditions. Eventually, the transcription factors (TF) of SDC2/4 were predicted based on the available dataset, and their involvement in septic inflammation was indirectly assessed. The GSE 5883 microarray gene-chip data profile was found to be suitable for analyzing lipopolysaccharide (LPS)-induced endothelial inflammatory injury. After analyzing the DEGs, 224 and 102 genes were found to be up- and down-regulated, respectively. All DEGs were involved in modulating receptor ligand and signaling receptor activator activities, cytokine receptor binding in biological processes, such as responses to lipopolysaccharide (LPS) and molecules from bacterial origins, positive regulation of cell adhesion, and other functions. In the case of localization of cellular components, most DEGs were enriched in cytoplasmic ribonucleoprotein granules, transcription regulator complex, and membrane rafts and were involved in the tumor necrosis factor (TNF) signaling and cytokine receptor interaction pathways. When compared with the control group, SDC4 gene expression in the LPS group increased by 4.5-fold, and comparable results were found in terms of SDC2 gene expression. From an experimental aspect, SDC4 was found to be markedly up-regulated with respect to mRNA and protein expression levels in response to inflammatory injury; moreover, down-regulation of SDC4 could severely exacerbate inflammatory responses either in in vivo or in vitro models. Altogether, SDC4, rather than SDC2, was found to be involved in LPS-induced sepsis-associated ALI.

## Introduction

It is clear that the pathogenesis of acute lung injury/acute respiratory distress syndrome (ALI/ARDS) leads to disturbances in lung homoeostasis due to virus- or bacteria-induced damage in the capillary endothelium or alveolar epithelium. This process leads to increased tissue permeability and tissue inflammation. Indirect ALI, mainly induced by sepsis and trauma, leads to changes in the microvascular endothelial cell structure and function and is extensively involved in the acute inflammatory response. This type of ALI accounts for high mortality in clinical settings [**?**]. It is known that the endothelium acts at the vascular alveolar septum, is more than a single nucleated layer, and can modulate a variety of functions, such as angiogenesis, blood clotting, inflammation, and expulsion of many molecules and proteins [**?**]. In normal states, endothelium positively facilitates the innate immune system to eliminate microbial invaders. In contrast, endothelial dysfunction can lead to a direct increase in vessel permeability thus leading to fluid leakage and edema formation. In addition to interrupting the inflammatory process initiated by neutrophile granulocytes due to the influence of adhesion of those accessory molecules with the vascular wall, intracellular signal transduction and inflammation cascades are disrupted [**?**]. To protect the innate immune response against overreacting, endothelial cells need to apply “brakes” to this process. Many studies have proven that endothelial dysfunction is connected with infiltration of neutrophils, expression of adhesion molecules, and activation of intracellular signaling pathways [**?, ?, ?**]. Furthermore, endothelial cells rely heavily on membrane surface syndecans (SDCs), which are membrane-attached proteoglycans, to repel the first pyemic “hit.” After all, any type of stimuli, such as cytokines, heparinases, and histamines can cause acceleration of glycocalyx degradation [**?**] and trigger inflammation development by interacting with heparin sulfate (HS) binding molecules circulating in the blood. At this point, it is urgent to elucidate the specific function of SDCs on the glycocalyx and to find therapeutic strategies to overcome sepsis-associated ALI.

To date, four types of SDCs have been found; however, only two of them, SDC2 and 4, are expressed on vascular endothelial cells. SDCs are membrane HS proteoglycan in heparan sulfate (HS) chains that attach to the ectodomain of the SDCs. Stimulation of SDCs can induce specific HS sulfation and dimerization, which uses SDCs’ special capabilities to interact with a large number of ligands and eventually span the duration of inflammatory responses and extent of activation of endothelial cells [**?**] [**?**] [**?**]. SDC4 is ubiquitous in many cells and participates in many different kinds of inflammatory responses. For instance, Godmann et al. found that SDC4 is involved in modulation of inflammatory arthritis through formation of a complex with interleukin 1 (IL-1) [**?**]. Estelle et al. reported that SDC4 can be expressed in islets and regulated by IL-1b. In lungs, CXCL8 and SDC4 can form a glycosaminoglycan-binding complex that induces inflammation [**?**]. Apart from these activities, many studies have demonstrated the involvement of SDC4 in various inflammatory conditions, such as rheumatoid arthritis [**?**], colitis [**?**], and cardiac inflammation [**?**]. As for SDC2, most related articles have primarily addressed angiogenesis, tumorigenesis, and metastasis aspects. In lungs, some studies have shed light on lung adenocarcinoma [**?**] and cancer [**?**]; nevertheless, researchers have rarely turned their attention toward the ALI field. SDC2 and 4 are structurally and evolutionary close and can likely inevitably have overlapping functions [**?**] [**?**]. Thus, a reason to evaluate their co-involvement in the ALI process exists.

In the present work, a bioinformatics technique was used to screen the DEGs in normal and LPS samples in human lung microvascular endothelial cells. A total of 326 genes were deferentially found and expressed. Among them, most genes were up-regulated and mainly participated in TNF signaling and cytokine receptor interaction pathways. Importantly, SDC4 expression increased several fold in these pathways. SDC2, however, maintained a comparably stable profile over time. To verify these results experimentally, LPS-induced in-vivo sepsis-associated ALI and endothelial injury in vitro models were established. mRNA and protein expression levels of SDC2 and 4 were examined in cell lysates and lung tissue, respectively, and the results were similar to those described above. At the end of the study, involvement of both candidate genes in the inflammatory process was tested again in an indirect manner by predicting the transcription factors and the knockdown of SDC4. Results showed that SDC4, rather than SDC2, could be transcribed by the nuclear factor kappa beta (NF-), the knockdown of which caused markedly worse inflammatory injury. Taken together, SDC4 (rather than SDC2) was found to be involved in sepsis-associated AL1; however, the exact mechanism and signaling pathway associated with this involvement need further study.

## Materials and methods

### Bioinformatics analysis and prediction

To predict and determine which isoform of SDC was involved in the endothelial damage on endotoxin attack, the Gene Expression Omnibus (GEO) database (http://www.ncbi.nlm.nih.gov/geo) was searched with the key words “LPS” and “endothelial cells,” and the appropriate dataset “GSE5883” was found, which was based on platform GPL570 with an Affymetrix Human Genome U133 Plus 2.0 Array. The study consisted of 24 samples of endothelial cells pretreated with LPS at 4, 8, and 24 h. According to our previous knowledge, an acute inflammatory response of ¡ 6 h was the longest time that could induce the changeable SDC expression. Four-hour data from GSE5883 were pooled and analyzed using the R-package (version 4.1.2). After normalization, a two-dimensional (2D) principal components analysis (PCA) was conducted to reveal the primary source of variability in the data. A gene expression matrix was then constructed with corresponding DEGs defined as the combination of p- value ¡ 0.01 and logFC ¿1.5. Several online websites and analytical tools, such as DAVID (https://david.ncifcrf.gov/), gene ontology GO; http://geneontology.org/), Kyoto Encyclopedia of Genomes and Genes (KEGG), and STRING (https://cn.string-db.org/,version 11.5) online websites in addition to cytoscape software were used to categorize and cluster the DEGs into specific modules based on their cellular function, cellular components, and biological processes. Moreover, for the prediction of possible signaling pathways consisting of those candidate DEGs, the KEGG website was used, and the output plot was depicted using R studio. To further investigate the expression level of SDCs in DEGs, SDCs were searched with the help of an R language in GSE5883 dataset, and the statistical results were analyzed using the t-test in R studio. Those molecules connected with SDC2/4 were also screened out.

### Experimental verification of isoform of SDCs

#### Construction of the experiment model

To verify the prediction derived from the above bioinformatics analysis, a sepsis-associated inflammatory injury model was constructed both in vivo and in vitro. Due to the possibility of human data involved in these models, the confirmation of ethics approval was achieved from the laboratory animal ethics committee of JXMC (JUMC2021-124) based on the fact that the experimental cells are immortalized human lung vascular endothelial cells (HLVECs), which were obtained from the Meisen Chinese Tissue Culture Collections. The cells were cultured normally in specific medium and activated with phosphate-buffered saline (PBS) as the control or LPS (LPS from Escherichia coli O55:B5) for 0, 6, 12, and 24 h with LPS doses ranging from 0 to 10 ug/ml. The concentration of IL-6 in the supernatant was examined using an enzyme-linked immunosorbent assay (ELISA) method. Additionally, the mRNA and protein expression levels of SDC2 and SDC4 in cell lysate were evaluated using a reverse transcriptase real time polymerase chain reaction (RT-qPCR) and western blotting. For the animal lung inflammatory model, LPS was administered intraperitoneally at 0, 6, 12, and 24 h time points with doses ranging from 0 to 30 ug/g. The concentrations of tumor necrosis alpha (TNF[U+237A]) and IL-6 in blood were measured using an ELISA to confirm the inflammatory status in the body after anesthesia. In vivo samples were obtained after mice were sacrificed by inhaling isoflurane. Hematoxylin and eosin (HE) staining and immunofluorescence examination of SDC2 and 4 expression in murine lungs at different time points was done. Meanwhile, the overall protein concentration in the bronchoalveolar lavage (BAL) was also determined.

### ELISA and immunofluorescence

After different procedures were completed, cell supernatant samples were collected from murine serum and BAL samples and centrifuged. Inflammatory biomarkers, such as TNF-a and IL-6, were examined using a precoated antibody ELISA from the manufacturer (Cat No. MAB2063-SP, DTA00D, RD Systems) following the manufacturer’s exact instructions. To determine which isoform of SDC was expressed in murine lung and the co-localization of SDC expression, lung samples of C57BL/6 mice from control group and the LPS group that received 20 *µ*g/g LPS were harvested after 24 h and concurrently embedded in 4 percent paraformaldehyde. Sections were then prepared, and double staining was finished with two primary antibodies, including anti-SDC4 antibodies (1:500, abcam ab286154) and anti-CD3 antibodies (1:500, abcam ab222783). Subsequently, a fluorescent-conjugated secondary antibody was added, and microscopic analysis was conducted to locate the molecular distribution of the fluorescent signals.

### RT-PCR experiment

Cells were harvested after washing twice with PBS, and RNA extraction was performed using the TRIZOL method. The concentration of overall RNA was determined using a nanodrop machine (ThermoFisher Scientific, USA). Using a commercial RNA reverse transcriptional Kit (PrimeScript RT Master Mix [Real Time], RR036A) and qPCR kit (TB Green Premix DimerEraser [Perfect Real Time], RR091A) provided by Takara Corporation (Takara Biomedical Technology [Beijing] Co., Ltd), quantitative real-time experiments were completed following the required protocols based on manufacturers’ instructions. Briefly, up to 1 *µ*g total RNA was reverse-transcribed to cDNA. The genetic assays were conducted in a 20 *µ*l reactive system containing Master Mix and 2 *µ*l of cDNA. The reaction parameters consisted of several specific conditions: (1) stage 1 at 95 ºC for 1 min, (2) stage 2 at 95 ºC for 5 s, and (3) 60 ºC for 1 min for 40 cycles. Subsequently, the relative gene expression levels of SDC2 and 4 were calculated in reference to the housekeeping gene, glyceraldehyde-phosphate dehydrogenase (GAPDH), accordingly. Apart from those ordinarily used primers, such as IL-6, IL-1, and vascular cell adhesion model 1 (VCAM-1), SDC2 and SDC4 primers used in this procedure are shown: (1) SDC2, F 5’-CTAGCAGCCGTCATTGC-3’, R 5’-TCCAAGGTCGTAGCTTCC-3’ and (2) SDC4, F5’-CGACGACGAAGATGCTG-3’, R 5’-CACCAAGGGCTCAATCAC-3’.

### Western blotting analysis

To determine the protein expression of SDC2/4 in vitro, cells were layered in a 6-well plate for both control and LPS groups. The cell lysates were prepared using RIPA lysis buffer containing several protease inhibitors. After examination of the overall protein concentration with a bicinchoninic acid (BCA) protein assay, 50 *µ*g lysate sample from each lane was added and separated using 12.5 percent sodium dodecyl gel electrophoresis (SDS-PAGE) gel for about 1 h. Subsequently, a 2-h membrane transfer experiment was completed, and 1 h blocking of membranes with milk plus 5 percent bovine serum albumin (BSA) was done. The primary antibodies were rabbit anti-SDC2 (1:1000, Abcam, ab205884) and rabbit anti-SDC4 (1:1000, Abcam, ab74139). Mouse anti-tubulin was used as an endogenous control gene. The membrane was incubated with these primary antibodies overnight in a 4 ºC refrigerator and then washed three times for 10 min after which horseradish peroxidase (HRP)-conjugated anti-rabbit second antibody (1:10000, Abcam, ab288151) was added to incubate at normal temperatures for 1 h. Finally, visualization was conducted on an enhanced bioluminescence instrument (Bio-Rad, USA). Each experiment was done in triplicate.

### Prediction of the TF of SDC4

To further explore the functional roles of SDC2/4 in the present experimental context, the GSEABase and clusterProfiler packages of R studio were used to calculate the possible transcription factors (TF) for SDC2 and 4 in the GSE5883 database to indirectly clarify their prospective involvement in inflammatory pathways. For SDC4, the prediction was based on the 87 DEGs used to retrieve all collected TF originating from C3 in the GSEA database (http://www.gsea-msigdb.org/gsea/msigdb/collections.jspC3), which incorporates all human TF and miRNA data. The summarized results were then constructed for SDC4 only and are illustrated in table below. The TF of SDC2, however, was deduced from all detected 17,690 genes on the gene chip due to the nonsignificant expression of its gene level, and a similar result was obtained as found for SDC4.

### Knock down of the SDC4 gene

In the in vitro model, small interfering RNA targeting the specific sequences of SDC4 was commercially designed by Tsingke (Shanghai, China). Meanwhile, the non-targeting siRNA was supplied as isotype control. Ten picomoles of SDC4 siRNAs (10 *µ*M) and 3 *µ*L Lipofectamine RNAiMAX (Invitrogen) were used as a complex to complete the transfection at 37ºC for 6 h; subsequently, the medium was changed to the complete medium for the remainder of the experiment. Verification of the knockdown was confirmed by the quantitative RT-PCR assay after 24 h of the transfection. As for the in vivo experiment, the pHBAAV-TIE-ZsGreen vector, developed by Hanheng Company, was used to construct the complete complex with the addition of sequence of Mir30-m-Sdc4 that targeted the specific sequence of SDC4 in endothelium to knockdown the mRNA expression of SDC4 (Mir30-m-Sdc4 sequence: AAG- GTATATTGCTGTTGACAGTGAGCGGATTCGAGAGACAGAGGTCATTAGT- GAAGCCACAGATGTAATGACCTCTGTCTCTCGAATCTGCCTACTGCCTCG). Subsequently, the PCR amplification and virus package were completed, and the HBAAV2/VEC-TIE-Mir30-m-Sdc4-ZsGreen virus was constructed with a titer of at least 1.1*1012 vg/mL. In the experimental group, the 65 *µ*l virus stock was transtracheally injected 30 days before construction of the model.

## Results

### identification of DEGs in GEO database

From the GSE5883 Expression profiling database, 8 samples at 4h studying time-point were divided into 4 control groups and LPS-treatment groups,respectively. With the criterion of Log2 Fold Change¿1.5 and p-value¡0.01 after uniformly standardization, 326 DEGs were found in total 17690 probes accordingly. By utilizing package ggplot of R studio(version 4.04), the sample data was extracted and the data matrix was built (Fig 1b).The expression profile was clustered and presented in 2 dimension PCA plot, which showed a obvious separation between control and LPS treatment samples, Moreover, the PC1 explained 19.6 percent variation of the transcriptomic profiles(Fig 1a).Among the contrast above, 326 DEGs altogether were found deferentially expressed and detected, interestingly, most of them were up-regulated(224 genes)due to the LPS stimulation(Fig 1d).The overall 326 DEGs are provided in Supplementary files S1.

**Fig 1.**
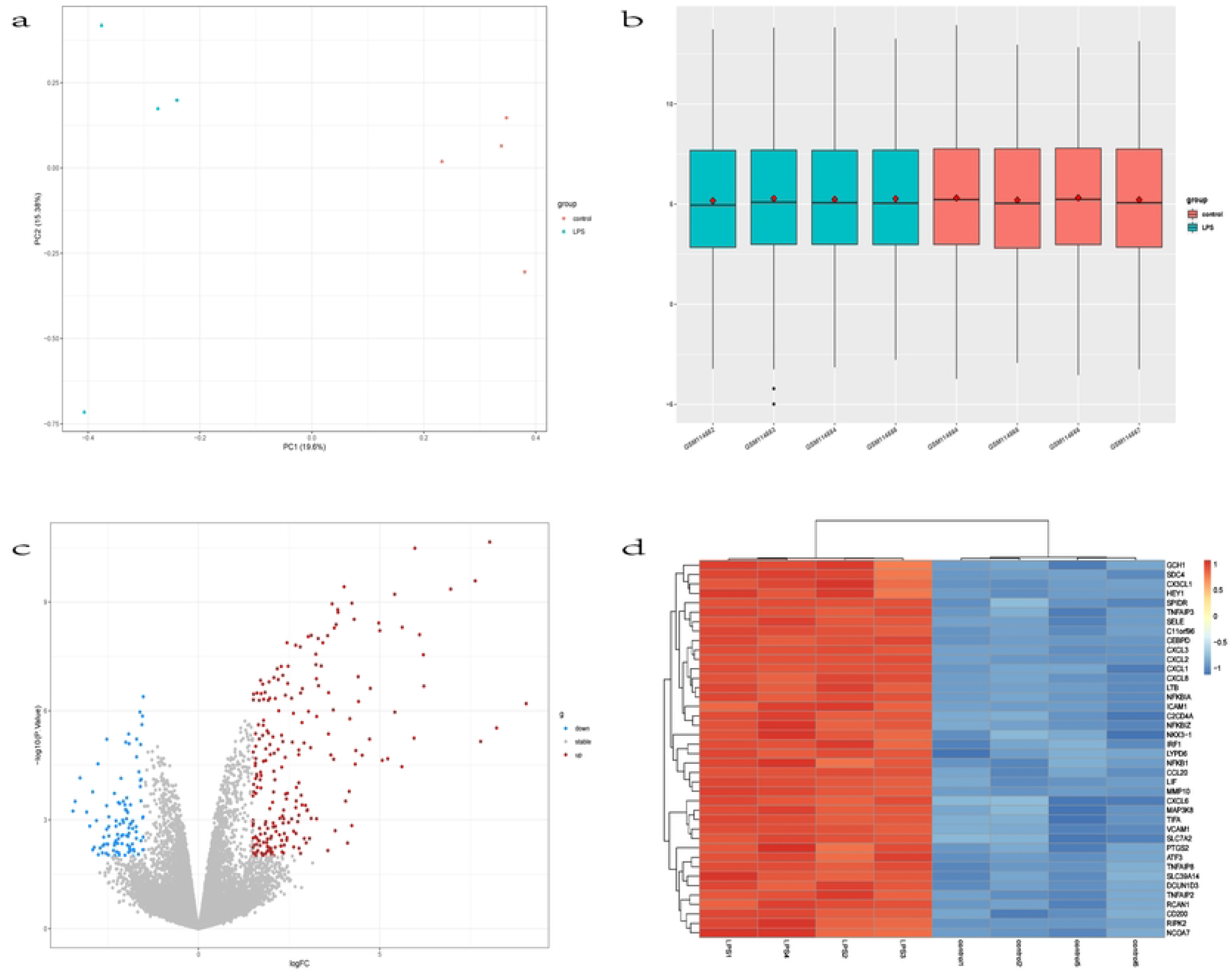
the plot presentation of outcomes of bioinformatics. (**a**) 2D PCA plot exhibition between 8 samples,the blue and red dots stands for LPS ans control samples,respectively. (**b**) Description of the general matrix expression.(**c**) the volcano plot of overall DEGs,the red and blue dots represent up-regulated and down-regulated genes,respectively.(**d**)heatmap of the first 40 DEG genes cited,the red and blue colors represent up-regulated and down-regulated genes,respectively.

### GO and KEGG analysis of DEGs

For further gene functional analysis, GO enrichment(available online website:http://geneontology.org/) and KEGG signaling pathway were conducted with 326 DEGs.The analyzed terms include biological process (BP), cellular components (CC), and molecular functions (MF).The top 5 significant GO terms for each enrichment were listed according to the gene count and p.value in figure(Fig 2a). It can be seen from the chart that genes in GO:0000785 had the lowest p.val, but the included gene numbers in the first 5 top terms were least.In other words, most numbers of genes were enriched in intracellular membrane-bounded organelle(270 genes),nucleus(195 genes),nuclear lumen(135 genes) and nucleoplasm(130 genes).In the molecular function part, 664 DEGs demonstrated th e most significant function of binding(GO:0005488) and protein binding(GO:0005515), those rest DEGs share relatively less participation in the cytokine activity(76 genes) and transcription regulator activity(26 genes). As for the biological process, the most popular process attended by DEGs was regulation of metabolic process, which counted 217 out of the whole number.besides, DEGs were also involved in the cellular response to cytokine stimuli,cellular response to organic substance and response to cytokine. From the online KEGG tool(Available online:http://www.genome.jp/kegg), functional signaling pathway enrichment of the top 20 pathways were pooled and depicted by R studio in figure(Fig 2b).It’s clear that the first and second important signaling pathways deployed by those DEGs were TNF signaling pathway and cytokine-cytokine receptor interaction pathway, which counted above 30 genes and with statistically obvious P.value(P¡0.00001). Subsequently, the NF-kappa B signaling, il-17 signaling and NOD-like receptor signaling were also effective candidates. As for the PPI cluster analysis of the DEGs analysed by cytoscape with a medium confidence(0.4), three clusters could be found in the supplementary file S2.

**Fig 2.**
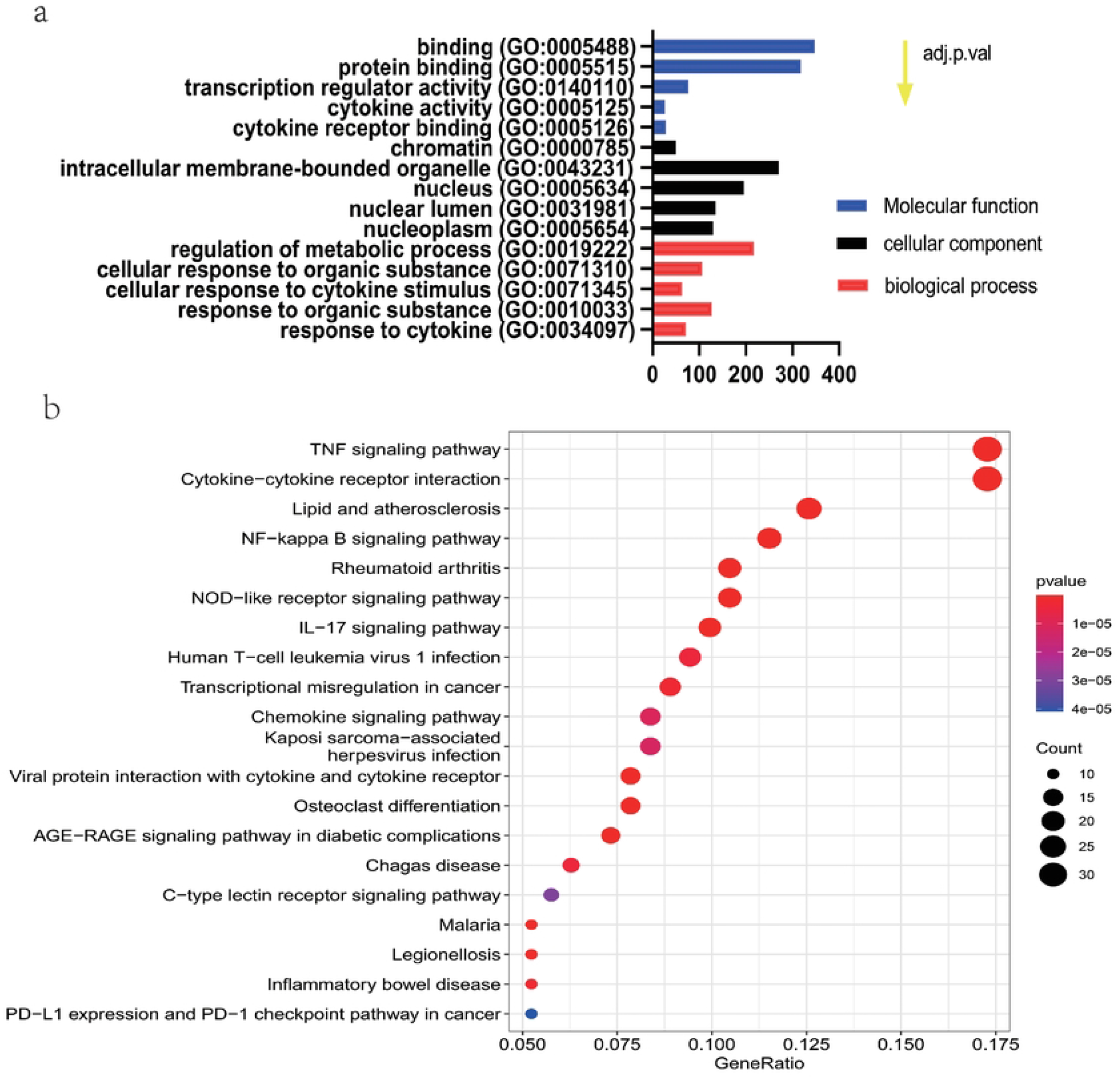
Gene ontology and Kyoto Encyclopedia of Genes and Genomes (GO and KEGG, respectively) enrichment of screened DEGs. The GO enrichment analysis was carried out on three categories: (1) molecular function, (2) cellular components, and (3) biological processes. Five GO terms are listed in Figure 2a according to the changes of adj. p.values and gene count. The most significant 20 presentations are illustrated with round bubbles; the rank was determined by the p-value and generation of signaling pathways shared by the DEGs in Figure 2b.

### SDCx genes expression in DEGs

After finishing those bioinformatics analysis above, the vague impression had been achieved about those DEGs. Frankly, most of them were involved in inflammatory reaction, and the LPS stimulation did induced remarkable inflammation inside the endothelial cells and inflammatory cascade signaling. However, in such an important reaction pathway, little is known about the role of SDCx and even which isoform of SDCx have the capacity to participate in those reactions. To this regard, the absolute quantification of SDCx were screened to verify the one, which has the capacity to locate through the menbrane structure of lung vescular endothelial cells and facilitate specific function. it truned out that apart from SDC1 and SDC3, both of which showed the average 5-6 quantitative expression(supplementary file S3a), SDC2 and SDC4 had 500-1500 quantitative expression level. Further, It could been seen that LPS-treatment stimulated above 4.5 fold increase of the SDC4 gene expression compared to control group(p¡0.001)(Fig 3a), nevertheless,the SDC2 expression showed a comparable result instead despite the fact that it’s also expressed on endothelial cells(Fig 3b). Using the cytoscape software for PPI network analysis, those molecules closely connected from SDC4 were also detected in cluster 1(the supplementary file S3b). CCL5, CXCL10,IL-1B, CXCL8 were four critical candidates.

**Fig 3.**
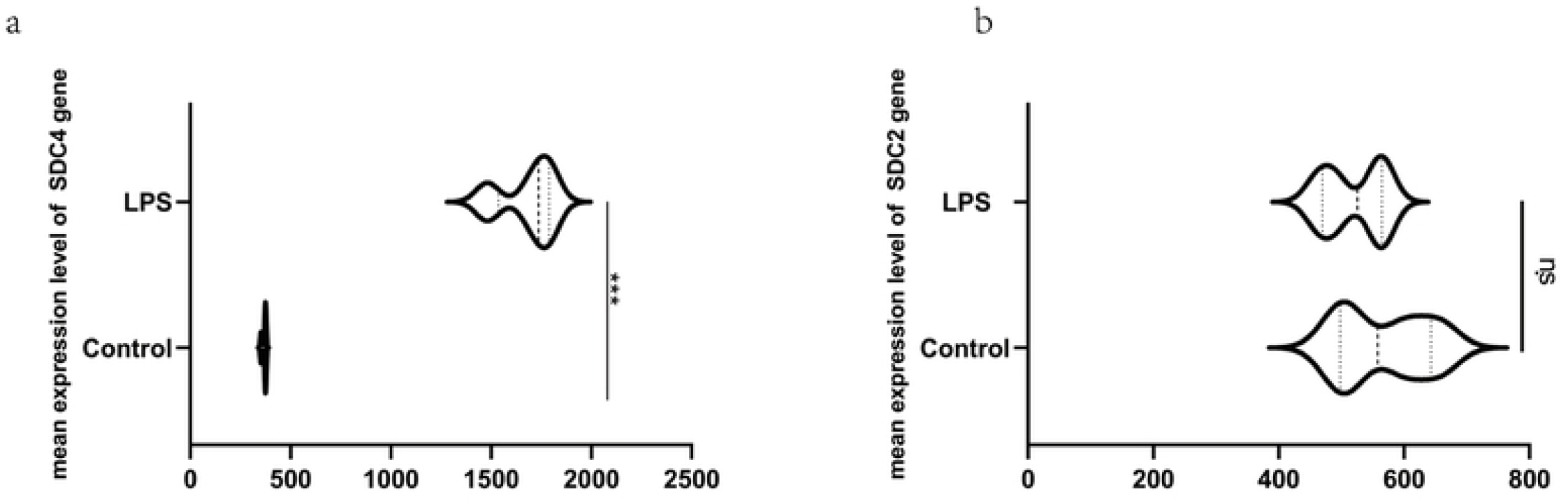
the primitive SDCx presentation in DEGs. (**a**) Compared to control group, LPS pretreatment induced dramatically high SDC4 expression. (**b**) Compared to control group, LPS pretreatment induced comparable SDC2 expression.n.s means not significant;*** represent p¡0.001.

### the verification of expression of SDCx in experiments

#### the build up of cellular and animal inflammatory injury model

The in vivo injury model was built up with 20ug/g LPS intraperitoneally injected in mouse, 0.5ml blood of each mouse was taken from Eye angular vein at 6hr,12hr and 24hr after build-up of model under anesthesia,respectively. Meanwhile, both of the lungs were taken after alveolar lavage was performed and the BAL fluid was collected. The result of HE staining of mouse lung tissue that directly reflect the extent of injury could be found in supplementary file S4. AS for the inflammatory indicators, TNF-a and IL-6 were found to be the extraordinary valuable molecules, the concentration of which markedly increased by the stimulation of LPS with a dose-dependent manner, especially at 20 and 30ug/g. the similar results were also found in BALF analysis(Fig 4a-d).In vitro inflammatory model, the statistically meaningful dose of LPS ranged from 0.5 to 10ug/ml, moreover, the time-course of the model was as early as 6hs after LPS administration to effectively induced the celluar inflammatory response witnessed by the IL-6 up-regulation in the Supernatant(Fig 4e-f).

**Fig 4.**
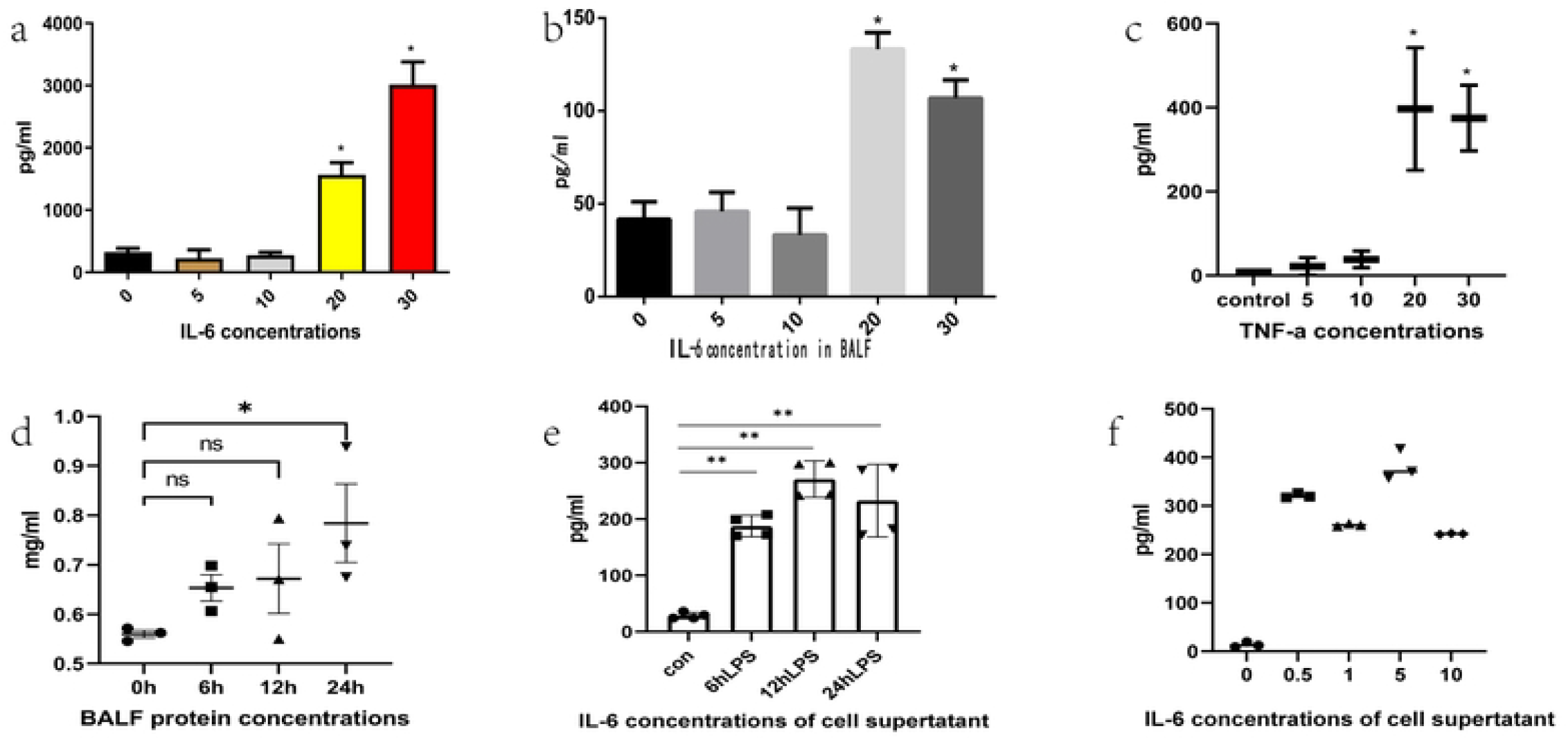
the inflammatory expression of experimental model on endothelial cell and C57BL/6 mouse. (**a**) the il-6 concentration in Lung homogenate of mouse when different doses LPS were administrated. (**b**)the il-6 concentration in BALF of mouse.(**c**) the TNF-a concentration in Lung homogenate of mouse. (**d**) the overall protein concentration of BALF.(**e-f**) the il-6 concentration in celluar supernatant when LPS was administered at several time points with different doses.BALF,bronchoalveolar lavage fluid,;n.s, not significant;***, p¡0.001.

#### the expression of SDCx in experimental models

As is well known that SDCx could be expressed by far more than endothelial cells, the lung tissue context is highly complex to be used for analysis of protein expression of interest, because the neutrophil and mononuclear macrophage could also have the capability to express these molecules under stimulus condition. For identify the subcellular location of SDC2/4 in lungs and even the co-localization with lung microvascular, double-staining with CD31(the endothelial cell marker) was conducted to clarify the anatomical localization. As can be seen from figure below(Fig 5), compared to control group(incubated with PBS) in vitro, LPS group(1ug/ml LPS administration for 6hr) had an marked increase both on mRNA and protein expression in terms of SDC4(P¡0.05). In contrast, SDC2 showed a comparable expression, although it’s also a membrane conserved proteoglycan(Fig 5a-c).In in vivo experiment, SDC2/4 both were identified in murine lungs no matter LPS induction or not, however, only SDC4 was found in co-staining image which showed a co-existence of SDC4 and CD31 in lung microvescular system(Fig 5d-e).

**Fig 5.**
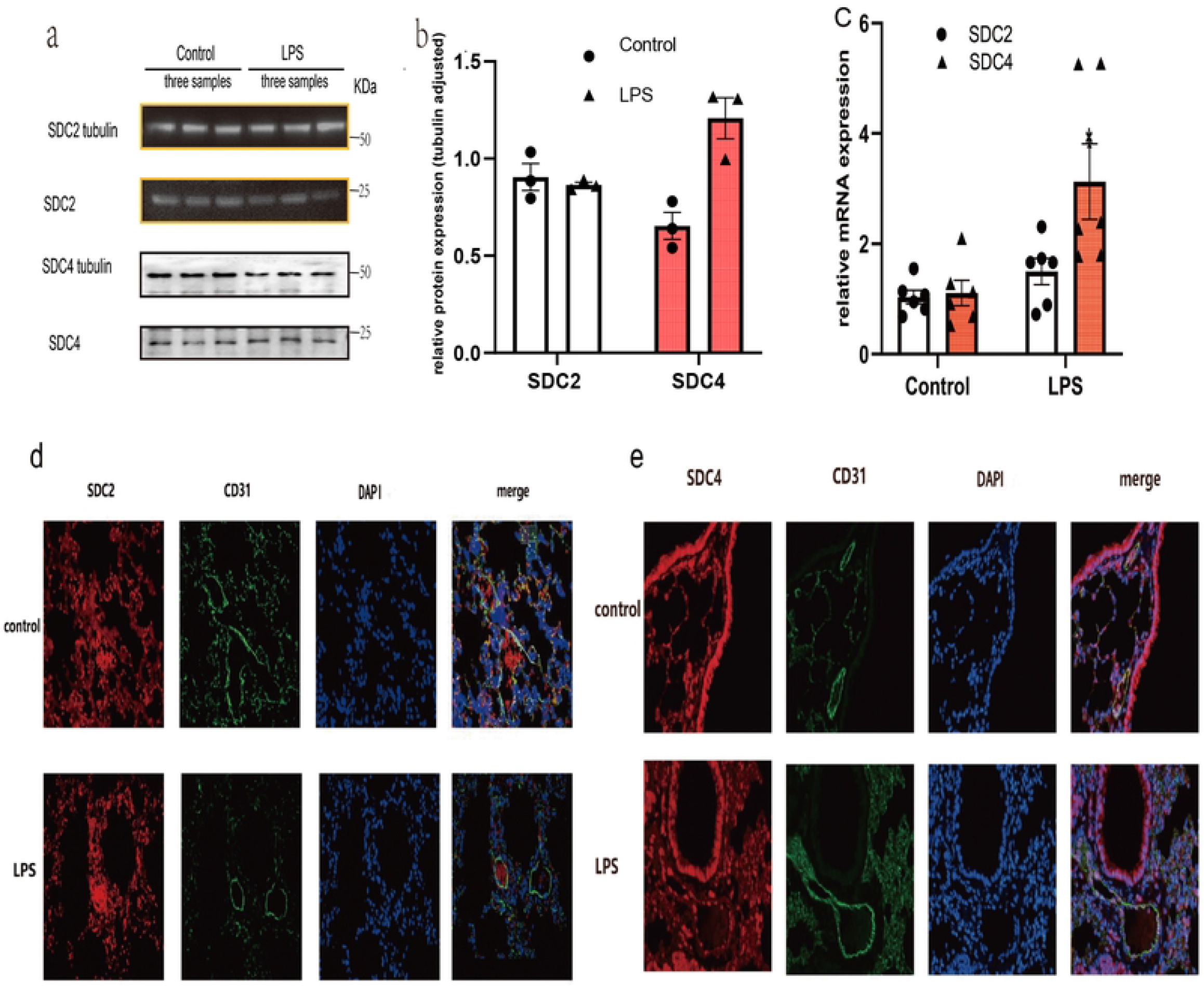
the SDC2/4 expression when inflammation appears in vivo and in vitro. (**a**) the relative SDC2/4 protein expression compared to -tubulin when 1ug/ml LPS was administered for 6hr, each group had triple biological repeats. (**b**)the statistical results of a.(**c**) the of SDC2 and SDC4 mRNA expression under LPS induction in vitro. (**d-e**) the SDC2/4 expression of inflammatory lungs in vivo for immunofluorescence exhibition and the co-staining of SDC2/4 with CD31. *,p¡0.05.

#### Regulation of acute inflammation by SDC4

As shown in (Fig 6), knockdown of SDC4 in vitro obviously led to an increase in the expression of three key inflammatory cytokines, IL-6 and -1 and VCAM-1, which are involved endothelial injury (Fig 6a–c); meanwhile, due to the overexpression of these cytokines induced by LPS in vivo, the HBAAV2/VEC-TIE-Mir30-m-Sdc4-ZsGreen virus caused a marked exacerbation in the lung injury exhibited in the HE pathological sections in fig 6d–f. These results show critically injured histology of lung tissue in KD group compared to LPS group, including the more thickened alveolar vascular barrier, more inflammatory cells, and hyaline membrane formation.

**Fig 6.**
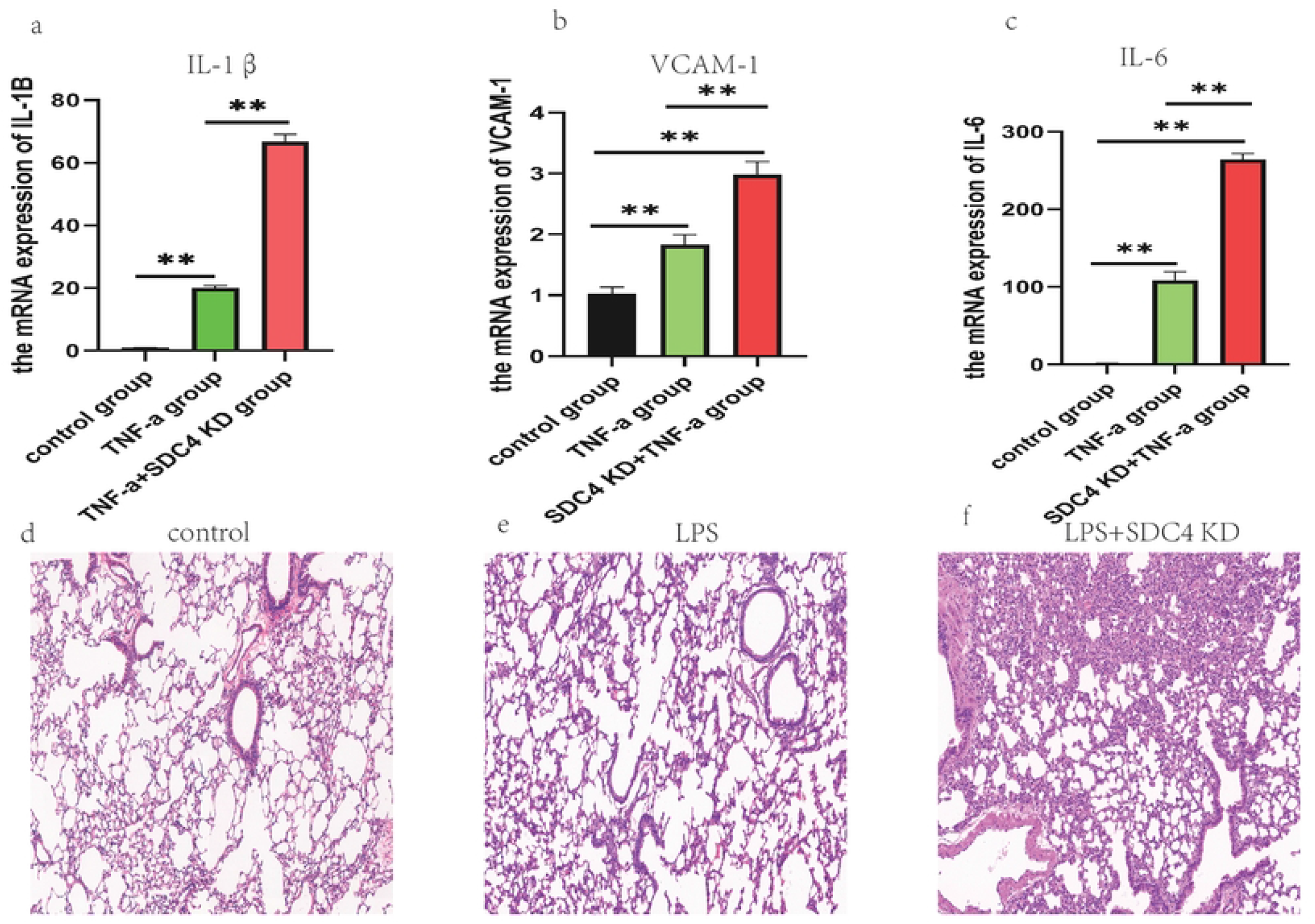
The inflammatory phenotype of in vivo and in vitro model mediated by the knockdown of SDC4. (**a-c**)the statistical results of the expression of three key inflammatory mediators in inflated endothelium. (**d-f**) the histology presentation of the HE pathological section of inflated lungs in mice. **, p¡0.01.

#### the transcription factors of SDC4

Through the pooled TF set from the result of prediction with the criterion of P¡ 0.05 in R studio, the predictive TFs for SDC2 and SDC4 were presented together in Table 1. Obviously, the TFs based on the pooled DEGs that could transcript SDC4 were NFKAPPAB, CREL and NFKAPPAB65.It’s easy to understand that NFKAPPAB and NFKAPPAB65 belong to NF-kappa B inflammatory signaling pathway, which is in charge of transcription of an array of downstream inflammatory biomolecules. CREL, encoded by gene REL, is also a NF-KB Subunit, facilitating the apoptosis, inflammation, the immune response, and oncogenic processes of most immune cells. As for SDC2, a different scenario was found in terms of its TF, which included ZNF512,ZNF512B and F10. According to explanation of genecards wedsite(https://www.genecards.org/), F10 encodes the vitamin K-dependent coagulation factor X of the blood coagulation cascade. ZNF512,ZNF512B encode a protein containing four putative zinc finger motifs, involving the phenotype of hair colour measurement, appendicular lean mass,et al, the whole material could be seen in supplementary files S5.

**Table 1.** The prediction of transcription factor of SDC2 and SDC4 by R Studio.

| SDC4 |  |  | SDC2 |  |
| --- | --- | --- | --- | --- |
| *predictedTF* | NFKAPPAB | 2.05E-20 | ZNF512 | 1.41E-12 |
| *andPvalue* | NFKAPPAB65 | 1.38E-13 | ZNF512B | 4.34E-15 |
|  | CREL | 5.14E-14 | F10 | 1.25E-11 |

SDC4, Syndecan 4; SDC2, Syndecan 2; TF, transcription factor

## Discussion

It is well known that the microvascular endothelial cell is the first “striking point” when sepsis-associated ALI (also called indirect ALI) is induced clinically. The SDCs and glycocalyx in the inner layer of vessels provide a significant contribution to the protection offered by endothelial cells and to chemotaxis of neutrophils, which are functionally completed through the binding of the heparin binding proteome. To elucidate the precise mechanism involved in the regulatory process between syndecan and the heparin-binding protein, the exact SDC isoform, its specific molecular function, and participating biological processes should be first clarified. In the present study, the SDC isoform of the lung microvascular endothelial cells, especially SDC2 and 4, which are involved in LPS-induced septic inflammation, were analyzed. The results show that SDC4 is the only isoform of SDC that is involved in LPS-induced sepsis-associated ALI; the possible involvement is related to inflammatory modulation. The definition of ALI refers to the early stages of acute lung injury, which usually occurs within first 72 h and which demonstrates systemic inflammation and a compromised homeostasis state between immune responses and insults to the immune system. ALI or ARDS usually appear when the balance is disrupted, and the inflammation has become an unstoppable event, such as sepsis, which is the most prevalent situation in clinical settings and is characterized by diffuse endothelial injury, coagulation imbalance, and hyperpermeability [18]. For indirect ALI, endothelial cell dysfunction is the leading cause that results in the development of ARDS and a consequence of pulmonary inflammation. With the increasing number of studies shedding light on the endothelial protection field, the glycocalyx and SCDs have attracted more and more attention due to their structural and functional roles. As reported earlier [19], the ectodomain of endothelial SCDs is degraded and shed under acute inflammatory conditions, a process that facilitates the downstream inflammatory cascade so that the concentrations of certain syndecans have become the biomarker of many deceases. For instance, in the case of the ongoing coronavirus 2019 (COVID-19) pandemic, the severity of pulmonary inflammation appears to be related to the degree of glycocalyx shedding in both infected children [20] and the elderly [21]. Therefore, some scholars even think about using both the endothelial cell and SDCs as promising therapeutic targets in clinical settings. In the lungs, the thickness of glycocalyx is of the utmost importance according to Schmidt’s study [3], which describes pulmonary SDC’s specific functions in dealing with acute inflammation. According to existing literature [22,23], SDC2 is mainly expressed in fibroblasts and epithelial and endothelial cells, and SDC4 is ubiquitous in almost every eukaryotic cell. In situations in which inflamed lungs are involved, SDC2 and 4 are inflammatory candidates to be further examined in terms of ALI/ARDS; furthermore, they should fulfill the principal criterion in which they are expressed during the early stage of inflammation and are involved in initiating the process. To match the acute phase of lung microvascular endothelial cells in the GEO database, GSE5883 was found to contain gene-chip data directly targeting lung microvascular endothelial cells and the acute phase at the 4 h after LPS induction. Through PCA analysis and normalized matrix distribution of the data profile, eight samples showed perfect non-variability and comparability. At the 4 h time points, 326 DEGs were found to be differentially expressed and detected according to the criterion of Log2 Fold Change ¿ 1.5 and p-value ¡ 0.01. Interestingly, most of the DEGs were upregulated (224 genes) after LPS stimulation. Subsequently, GO and KEGG enrichment were conducted to characterize these DEGs. In the present bioinformatics analysis, most DEGs were involved in the TNF and cytokine receptor signaling pathways, which is compatible with the experimental background. As for GO enrichment, the results indicate that DEGs had the capacity to bind other molecules (GO:0005488) or proteins (GO:0005515). The biological processes included regulation of metabolic process, cellular responses to cytokine stimuli and organic substances, and others. Furthermore, these DEGS were primarily located in the intracellular membrane-bound organelles or nucleus. These results indicate that endothelial cells mainly use intracellular signal transduction or transcribe most of inflammatory molecules when inflammation is present. As described in Fu and colleagues’ study [24], which used an integral bioinformatics analysis to determine the key genes involved in sepsis, they found 619 common overlapping genes due to different enrollment criterion than used in our study. Nevertheless, their results shared a similar conclusion to ours, namely, protein binding based on GO function, the cytoplasm and integral components of membrane cellular components, and the same pathway, such as the TNF signaling pathway. As for the function of extracellular matrix proteins, these proteins are not a major phenotype with which to be concerned. We could not draw the conclusion that those membrane-attached proteoglycans have only minor importance for modulating the initiation of ALI. Among the total 17690 genes, mRNA expression of SDC1-4 could be measured when compared with the control group, but only SDC2/4 showed an absolute quantitative rise in gene expression of more than 100-fold than SDC1/3 (details can be seen in supplementary file S3). A 4.5-fold statistical change was seen in SDC4 versus SDC2 expression, which showed a comparable result, thus emphasizing the important role of SDC4 in the ALI pathway. Moreover, the PPI molecules associated with SDC4 included CCL5, CXCL 8 and 10, and IL-1B, all of which had a statistically significantly meaning in the present analysis. Among them, CCL5(P13501), CXCL10(P10145), and CXCL8(P10145) belong to chemotactic factors characterized by the capability to bind their respective receptors and further induce the migration of inflammatory or immune cells to the infected sites (explanation from uniprot), which are all cellular response to organic cyclic compound and inflammatory response. IL-1B(P01584) is a potent proinflammatory cytokine possessing cytokine activity. Most important, all of these molecules share a common characteristic in that they bind to glycosaminoglycan or the HS chains of SDC isoforms or undergo heparin binding. Based on a search of PubMed, SDC isoforms are prone to bind CCL5 [25], CXCL10 [26], IL-1B [27], and CXCL8 [28] in diverse context in addition to inflammation to modulate intracellular signaling transduction or function as co-receptors to facilitate other pathways. As for the specific mechanisms, these are not addressed in this study. Once again, the prediction of TF that could possibly transcribe SDC4 was necessary to enhance the strength of evidence, it turned out that in this septic context SDC4 could be transcribed by NF- and its subfamily members. In experiments, this possibility of NF- to SDC4 transcription has previously been verified in the inflammation and cancer fields [29,30]. Altogether, these facts above at least illustrated directly or indirectly that SDC4 is a potential syndecan in ALI. To verify in vivo and in vitro expression of SDC2 and SDC4 experimentally, the models induced by LPS were used. First, an in vivo sepsis model was tested by determining pathological and morphologic changes of lung tissue, the wet/dry ratios, and injury scores, which verified the drastic changes in LPS group at 24 h after LPS administration. Based on this model, TNF-a and IL-6 levels were found to have significantly increased. This evidence was enough to validate the construction of an in vivo model. SDC4, in contrast to SDC2, was co-stained with CD31, which is a blood marker, in both the control and LPS groups. As for the differential expression between groups no need to clarify our findings exists at present. Second, the IL-6 concentration was upregulated at a broad volume range since LPS was administrated for 6 h, 1 ug/ml LPS resulted in the remarkable elevation of mRNA and protein expression of SDC4, nevertheless, the expression of SDC2 showed no changes. In this part, LPS was chosen as a stimulation because LPS has similar capability as TNF-a and IL-1B to induce inflammation, and the selected volume was based on our pilot study. SDC4, due to the chondroitin sulfate (CS) side chains in the glycosaminoglycan, has been fully studied in osteoarthritis [31–33]. With the help of HS chains, SDC4 is involved in the regulation of cancer [25,34] or other diseases. In the ALI/ARDS fields, the importance of SDC4 has increasingly been increasing revealed. Tram and colleagues [35] reported SDC3 and 4 were all up-regulated by inflammatory stimuli, but the experimental cells used were human umbilical vein endothelial cells, which are not relevant in lung studies. Arif et al. [36] found SDC4 could modulate the Keap1-Nrf2 antioxidant pathway in the lung to cause a reduction in inflammation when administrated systemically. This systemic administration indirectly illustrates the interaction between SDC4 and lung microvascular endothelial cells. Even some research has reported that SDC4 may substitute for SDC1 in ALI. It can be concluded that SDC4 is the only available glycosaminoglycan involved in the pulmonary inflammation. Several limitations should be discussed in terms of the present study. First, only one GSE database was adopted for analyses and may not be sufficient, but the examples and matrix distribution were perfect, and the conclusion was compatible with other studies. Second, TF is predicted in a limited territory, the purpose was to support the prediction, the standard verification needs further double luciferase validation test. Third, based on our results, the peak of inflammatory state was found to occur 24 h after LPS induction, while SDC4 was found to be expressed at a relatively early phase (6 h). If SDC4’s function elapse in later period of time was not included in this study, an explanation for this finding is warranted in forthcoming studies. Last but not least, the in vivo SDC4 protein expression level in endothelium could not be obtained due to the limitations in the technique.

## Conclusion

With the help of a bioinformatic analysis and basal experiments, SDC4 was screened in the GEO dataset and found to be the only inflammatory regulator of interest; moreover, SDC4 involvement was once again verified to be protective when examining lung inflammatory injury.

## Disclosure

The author reports no conflicts of interest in this work.

## Acknowledgments

We thank the invaluable assistance of professor Lina Yu (Department of Anesthesiology, the Second Affiliated Hospital of Medical College of Zhejiang University) and Dr Gaojian Wang (Department of Anesthesiology, the Run Run Shaw Hospital of Medical College of Zhejiang University) during the whole experiment.

